# Omics Pipe: A Computational Framework for Reproducible Multi-Omics Data Analysis

**DOI:** 10.1101/008383

**Authors:** Kathleen M. Fisch, Tobias Meissner, Louis Gioia, Jean-Christophe Ducom, Tristan M. Carland, Salvatore Loguercio, Andrew I. Su

## Abstract

Omics Pipe (https://bitbucket.org/sulab/omics_pipe) is a computational platform that automates multi-omics data analysis pipelines on high performance compute clusters and in the cloud. It supports best practice published pipelines for RNA-seq, miRNA-seq, Exome-seq, Whole Genome sequencing, ChIP-seq analyses and automatic processing of data from The Cancer Genome Atlas. Omics Pipe provides researchers with a tool for reproducible, open source and extensible next generation sequencing analysis.

## Main

Next generation sequencing (NGS) has presented researchers with the opportunity to collect large amounts of sequencing data^1^, which has accelerated the pace of genomic research with applications to personalized medicine and diagnostics^2^. This has resulted in the development of a large number of computational tools and analysis pipelines, necessitating the creation of best practices and reproducible integrative analysis frameworks^3^. The Nature Protocols journal has attempted to create one solution to establish best practices, by publishing step-by-step directions for well-established NGS analysis pipelines. In addition, The Broad Institute has outlined best practices for variant calling using the Genome Analysis Toolkit (GATK)^4^, ENCODE has recommended experiment guidelines and best practices for personalized genomic medicine are emerging^5^. Several automated pipelines have been developed to tie together individual software tools^2^, although many of these tools focus on a single NGS platform, require computational expertise, require commercial licenses and/or are poorly documented3.

To address these issues, we developed Omics Pipe (https://bitbucket.org/sulab/omics_pipe), an open-source, modular computational platform that automates best practice multi-omics data analysis pipelines with built-in version control for reproducibility (Figure 1). It currently supports two RNA sequencing (RNA-seq) pipelines ^6,7^, variant calling from whole exome sequencing (WES) and whole genome sequencing (WGS) based on GATK ^4^, two ChIP-seq pipelines ^8,9^ and custom RNA-seq pipelines for personalized cancer genomic medicine reporting and analysis of The Cancer Genome Atlas (TCGA) datasets^10^ (Supplementary Figure 1).

**Figure 1.**
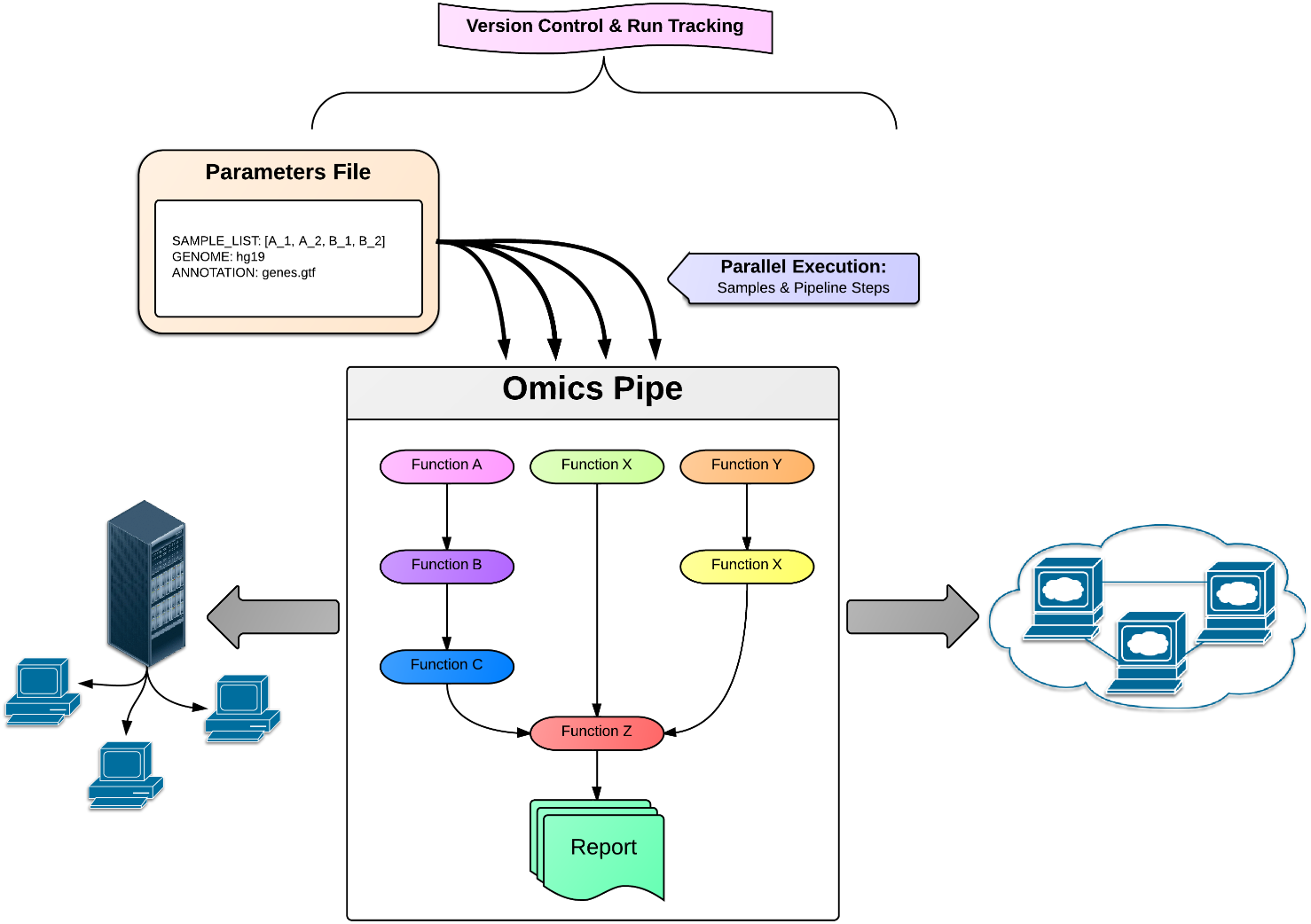
Schematic diagram of Omics Pipe.

Omics Pipe is distributed as a lightweight Python package that can be installed on a compute cluster, a local installation or in the cloud. It is hosted on PyPi for direct download for local and cluster installation, and it is also hosted as an Amazon Machine Image (AMI) in Amazon Web Services (AWS) Elastic Compute Cloud (EC2). The AWS distribution of Omics Pipe runs on MIT’s StarCluster (http://star.mit.edu/). The AMI is preconfigured with all of the required software dependencies to run best practice next generation sequencing analyses, providing a valuable computational resource to the scientific community. A Docker container (https://www.docker.com/) is provided to configure and boot up StarCluster with the preconfigured Omics Pipe AMI and preinstalled third-party software dependencies. Detailed tutorials (http://pythonhosted.org/omics_pipe/), documentation and source code are hosted in an open source repository that will allow community contribution to the source code as well as transparency for reproducibility and accuracy (https://bitbucket.org/sulab/omics_pipe).

The Omics Pipe framework is modular, which allows researchers to easily and efficiently add new analysis tools with scripts in the form of Python modules that can then be used to assemble a new analysis pipeline. Omics Pipe depends upon the Python package Ruffus^11^ to pipeline the various analysis modules together into a parallel, automated pipeline. This also allows for the restarting of only the steps in the pipeline that need updating in the event of an error. In addition, Sumatra ^12^ is built into the Omics Pipe framework, which provides version control for each run of the pipeline, increasing the reproducibility and documentation of the analyses.

Omics Pipe automatically submits, controls and monitors jobs on a Distributed Resource Management system, such as a compute cluster, MIT’s StarCluster or Grid computing infrastructure. This allows samples and steps in the pipeline to be executed in parallel in a computationally efficient, distributed fashion, without the need to individually schedule and monitor individual jobs. For the RNA-seq analysis pipelines, an analysis summary report will be generated as an HTML report using the R package knitr^13^. The summary report provides quality control metrics and visualizations of the results for each sample to enable researchers to quickly and easily interpret the results of the pipeline.

One advantage of a robust and simple to use pipeline is the ability to easily reanalyze existing data sets using the most recent algorithms and annotations. To illustrate this point, we used Omics Pipe to reanalyze a subset of the breast invasive carcinoma RNA-seq dataset (N = 100) paired tumor-normal samples generated by the TCGA Research Network (http://cancergenome.nih.gov/) using the count-based differential expression analysis best practice protocol^6^ and updated UCSC RefSeq annotations (V57). Omics Pipe automatically downloaded the desired TCGA samples and ran the selected pipeline on a high throughput compute cluster. We performed paired differential expression analysis, signaling pathway impact analysis and consensus clustering analysis using the Bioconductor packages edgeR^14^, SPIA^15^ and ConsensusClusterPlus^16^, respectively. We then compared the results of the reanalysis to the original TCGA RNAseq V2 Workflow (UCSC RefSeq General Annotation Format 2011), by downloading the raw counts for the same samples from TCGA and performing the analyses described above.

The updated UCSC RefSeq V57 annotations contained 3,475 additional genes compared to the UCSC RefSeq General Annotation Format from 2011 used to originally analyze the TCGA data (Figure 2a). The reanalysis of the TCGA breast invasive carcinoma samples using Omics Pipe revealed 761 differentially expressed (DE) genes compared to the original TCGA analysis, which resulted in 410 DE genes (Supplementary Tables 1 & 2). There were 394 DE genes shared between the two analyses (Figure 2b). In the reanalyzed dataset, 367 DE genes were unique, 14 of which were due to new annotations. One of the newly annotated DE genes, DSCAM-AS1, was upregulated 256x in tumor versus normal samples, which has been implicated in the malignant progression of carcinomas by an estrogen-independent mechanism^17^.

**Figure 2.**
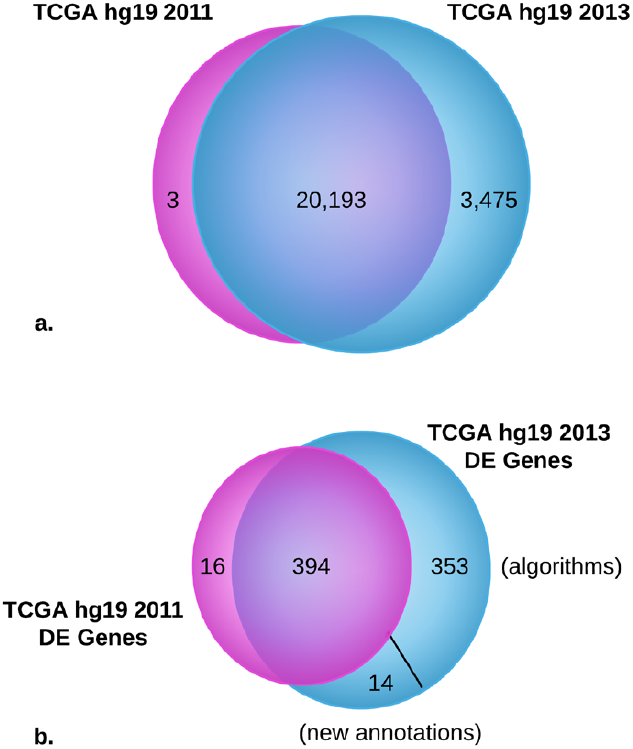
Comparison of the number of genes annotated in two different UCSC RefSeq releases and the number of differentially expressed (DE) genes identified by different algorithms and annotations **(a)** Venn diagram of the number of genes annotated in the UCSC RefSeq hg19 2011 Generic Annotation File and the UCSC RefSeq hg19 2013 annotation (Release 57) **(b)** Venn diagram of the comparison of the number of DE genes identified between raw counts generated with the TCGA UNC V2 RNA-seq Workflow using the UCSC RefSeq hg19 2011 Generic Annotation File and raw counts generated with the count-based pipeline in Omics Pipe using the UCSC RefSeq hg19 2013 annotation (Release 57).

Consensus clustering of the original TCGA counts resulted in four clusters, with each cluster containing both tumor and normal samples (Supplementary Figures 2 & 3). Consensus clustering of the reanalyzed counts resulted in 10 clusters, with tumor and normal samples clustering separately, with the exception of one normal sample clustering with two tumor samples in Cluster 7 (Supplementary Figures 2 & 3). These results indicate that the addition of the 3,475 genes in the new annotation provides additional information to improve the separation of tumor and normal samples. Twenty significantly dysregulated pathways were identified from the DE genes from the original TCGA counts, and 29 significantly dysregulated pathways were identified in the reanalyzed dataset, 11 of which were new pathways primarily related to RNA polymerase activity (Supplementary Tables 3 & 4). The reanalysis of the TCGA data using a best practice pipeline and updated annotations demonstrates the utility of Omics Pipe as a tool for conducting reproducible NGS analyses that can lead to novel biological insights.

In conclusion, Omics Pipe is an automated and reproducible computational framework that can be used to efficiently analyze newly generated data or to reanalyze publically available data, such as TCGA. It currently supports several best practice pipelines for RNA-seq, WES, WGS and ChIP-seq. This list of pipelines will continue to be updated, and we invite the broader community to participate in the continued development of Omics Pipe through our open source code repository. Pull requests for new components and new pipelines will be properly reviewed. In addition, the built-in version control system allows for the reproducibility of analyses performed within the Omics Pipe framework, which is important as new versions of software tools and annotation are released. It can be easily extended as new tools become available, and it can be implemented on a local machine, a computer cluster or the cloud. The goal of Omics Pipe is to democratize NGS analysis by dramatically increasing the accessibility and reproducibility of best practice computational pipelines, which will enable researchers to generate biologically meaningful and interpretable results.

## Methods

Methods and associated references are available in the online version of the paper.

## ONLINE METHODS

### The Omics Pipe Framework

Omics Pipe is a Python package that pipelines shell scripts into an automated, version controlled, parallelized pipeline of steps based on the Python package Ruffus11 for running the pipeline steps, Sumatra12 for version control and run tracking, and Python DRMAA (https://github.com/drmaa-python) for distributed computing. Omics Pipe is distributed as a standalone Python package for installation on a local cluster already containing the third-party software dependencies and reference databases. Omics Pipe is also distributed as an Amazon Machine Image (AMI) in Amazon Web Services (AWS) Elastic Compute Cloud (EC2) that contains all necessary third-party software dependencies and databases. The AWS distribution of Omics Pipe runs on MIT’s StarCluster (http://star.mit.edu/). A Docker container (https://www.docker.com/) is provided to configure and boot up StarCluster with the preconfigured Omics Pipe AMI.

Omics Pipe requires the user to specify which supported pipeline to execute at the command line or specify the path to a custom Python script containing a custom pipeline. The user also must supply a parameter file in YAML format to feed in relevant parameters for running the pipeline, including the command line options for each of the tools and other customizable settings. All of the parameters have default values to enable the user to run the supported pipelines with minimal start up time. More advanced users can customize every option possible from each of the pipelined tools.

Omics Pipe can be extended by the user to create custom pipelines from built-in modules and by creating modules for new tools. It is language agnostic, so existing shell scripts written in any programming language can be included as an Omics Pipe module. Omics Pipe executes the shell scripts on the cluster or in the cloud using DRMAA to allocate resources and manage job execution. Omics Pipe checks that the job finished successfully and creates a flag file upon successful completion, allowing the user to rerun only out of date steps in the pipeline. Ruffus11 provides functionality for parallel execution of pipeline steps. Each time Omics Pipe is executed, Sumatra12 creates a database entry to log the specifics of the run, including the parameters, input files, output files and software versions for version control and run tracking.

## Supported Best Practice Pipelines

Omics Pipe currently supports six published best practice pipelines. These include two RNA sequencing (RNA-seq) pipelines for both mRNA and miRNA6,7, variant calling from whole exome sequencing (WES) and whole genome sequencing (WGS) based on GATK4, and two ChIP-seq pipelines8,9. It also includes custom RNA-seq pipelines for personalized cancer genomic medicine reporting and analysis of The Cancer Genome Atlas (TCGA) datasets10. The steps in each method have been adapted exactly as described in the associated publications, allowing the user to implement these methods on their own datasets. The command-line options for each tool in each pipeline are exposed to the user in the parameters file.

## Using Omics Pipe to Automatically Analyze TCGA data

We used Omics Pipe to reanalyze 100 paired tumor/normal samples from 50 patients in the TCGA breast invasive carcinoma dataset to demonstrate its utility for efficiently processing samples using best practice pipelines. We automatically downloaded the raw RNA-seq fastq files and processed the files using the count-based differential expression analysis best practice protocol6 to quantify gene expression.

Briefly, sequencing reads were aligned to the human genome (hg19) using the STAR aligner18. Gene expression quantification was performed at the exon level using the htseq-count function within the Python HTSeq analysis package19 with UCSC RefSeq hg19 annotation (Release 57). Nonspecific filtering was applied to the raw count data and the 50% most variable genes were used in the differential expression analysis after TMM normalization. Differential gene expression was performed using a paired design matrix with the Bioconductor package edgeR14. Genes with a False Discovery Rate (FDR) < 0.01 and log2(FoldChange) > |2| were considered differentially expressed.

We also downloaded the raw count files generated from the TCGA UNC V2 RNA-seq Workflow for the same 100 samples. These counts were generated using the UNC V2 RNA-seq

Workflow and were based on the UCSC RefSeq hg19 Generic Annotation File from June 2011. We performed differential expression analysis of these raw counts as described above.

## Identification of Novel Genes, Pathways and Clustering in TCGA Breast Invasive Carcinoma

We compared the differentially expressed genes in the reanalysis of the TCGA dataset using Omics Pipe to the raw counts processed by TCGA to assess the utility of rerunning previous analyses with updated gene annotations and algorithms. We updated the gene identifiers provided with the original raw count data using the R package mygene.R (https://bitbucket.org/sulab/mygene.r) and we extracted newly annotated differentially expressed genes identified in the reanalyzed dataset.

We performed a Signaling Pathway Impact Analysis (SPIA) to identify significantly dysregulated pathways with the Bioconductor packages SPIA^15^ and Graphite^21^ based on the Biocarta, KEGG, NCI and Reactome databases. We performed this analysis once on each dataset, using the differentially expressed genes in each dataset as input, and setting the background genes to all genes included in the differential expression analysis for each dataset.

We identified patterns of relationships among the samples in each dataset using the Bioconductor package ConsensusClusterPlus16 with 80% resampling from 2 to 20 clusters and 1,000 iterations of hierarchical clustering based on a Pearson correlation distance metric. We compared the differentially expressed gene sets, pathways and clusters between the previously published results and the current analysis using updated annotations to identify novel differentially expressed genes and pathways relevant to breast cancer.

## Author contributions

K.F. designed the research, developed the pipeline framework, coded the pipeline scripts, performed and validated the results and wrote the manuscript. T.M. participated in designing the pipeline framework, coded the pipeline scripts, participated in developing the AWS distribution and wrote the manuscript. L.G. developed the AWS distribution. J.D. participated in designing the pipeline framework and provided computational support and guidance. T.C. participated in coding the pipeline scripts. S.L. participated in coding the pipeline scripts. A.S. designed and supervised the research and participated in writing the manuscript.

## Acknowledgements

This work was supported by the National Center for Advancing Translational Sciences (Grant UL1TR001114), the National Cancer Institute (Grant CA92577), the National Institute on Alcohol Abuse and Alcoholism (Grants AA007456, AA013525), the National Institute on Drug Abuse (Grant DA030976), and by a fellowship from the National Foundation for Cancer Research.

## Supplementary Tables & Figures

**Supplementary Figure 1.** Pre-built best practice pipelines and the third party software tools supported by Omics Pipe. Users can easily create custom pipelines from the existing modules and they can create new modules supporting additional third party software tools.

**Supplementary Figure 2.** Consensus clustering analysis of the TCGA breast invasive carcinoma paired tumor-normal samples performed with the reanalyzed count data (a)-(d) and the original raw counts downloaded from TCGA (e)-(h) for cluster sizes of k=2, k=3, k=4 and k=10. The blue and white heat map displays sample consensus.

**Supplementary Figure 3.** Measurements of consensus for different cluster sizes (k) from the consensus clustering analysis on the reanalyzed (a)-(c) and original counts (d)-(f) from the TCGA paired tumor-normal breast invasive carcinoma samples. The empirical cumulative distribution (CDF) plots (a) and (d) indicate at which k the shape of the curve approaches the ideal step function. Plots (b) and (e) depict the area under the two CDF curves. Item consensus plots (c) and (f) demonstrate the mean consensus of each sample with all other samples in a particular cluster (represented by color).

**Supplementary Table 1.** Differentially expressed genes (FDR < 0.01 and log2(FoldChange) > |2|) resulting from the reanalysis of the paired tumor-normal samples from the TCGA breast invasive carcinoma dataset using the count-based best practice pipeline with UCSC RefSeq V57 annotation in Omics Pipe.

**Supplementary Table 2.** Differentially expressed genes (FDR < 0.01 and log2(FoldChange) > |2|) resulting from differential expression analysis of the original raw counts from the paired tumor-normal samples from the TCGA breast invasive carcinoma dataset.

**Supplementary Table 3.** Significantly dysregulated pathways in the reanalysis of the paired tumor-normal samples from the TCGA breast invasive carcinoma dataset using the count-based best practice pipeline with UCSC RefSeq V57 annotation in Omics Pipe.

**Supplementary Table 4.** Significantly dysregulated pathways from differential expression analysis of the original raw counts from the paired tumor-normal samples from the TCGA breast invasive carcinoma dataset.

